# CatPred: A comprehensive framework for deep learning in vitro enzyme kinetic parameters *k_cat_*, *K_m_* and *K_i_*

**DOI:** 10.1101/2024.03.10.584340

**Authors:** Veda Sheersh Boorla, Costas D. Maranas

## Abstract

Quantification of enzymatic activities still heavily relies on experimental assays, which can be expensive and time-consuming. Therefore, methods that enable accurate predictions of enzyme activity can serve as effective digital twins. A few recent studies have shown the possibility of training machine learning (ML) models for predicting the enzyme turnover numbers (*k_cat_*) and Michaelis constants (*K_m_*) using only features derived from enzyme sequences and substrate chemical topologies by training on *in vitro* measurements. However, several challenges remain such as lack of standardized training datasets, evaluation of predictive performance on out-of-distribution examples, and model uncertainty quantification. Here, we introduce CatPred, a comprehensive framework for ML prediction of *in vitro* enzyme kinetics. We explored different learning architectures and feature representations for enzymes including those utilizing pretrained protein language model features and pretrained three-dimensional structural features. We systematically evaluate the performance of trained models for predicting *k_cat_*, *K_m_*, and inhibition constants (*K_i_*) of enzymatic reactions on held-out test sets with a special emphasis on out-of-distribution test samples (corresponding to enzyme sequences dissimilar from those encountered during training). CatPred assumes a probabilistic regression approach offering query-specific standard deviation and mean value predictions. Results on unseen data confirm that accuracy in enzyme parameter predictions made by CatPred positively correlate with lower predicted variances. Incorporating pre-trained language model features is found to be enabling for achieving robust performance on out-of-distribution samples. Test evaluations on both held-out and out-of-distribution test datasets confirm that CatPred performs at least competitively with existing methods while simultaneously offering robust uncertainty quantification. CatPred offers wider scope and larger data coverage (∼23k, 41k, 12k data-points respectively for *k_cat_, K_m_ and K_i_*). A web-resource to use the trained models is made available at: https://tiny.cc/catpred

## Introduction

Continued advances in genomics and metagenomics tools have spearheaded an unprecedented pace in the discovery of new genetic sequences^1^. While the growth of newly deposited genetic sequences within genomic databases^2^ maintain an exponential rate, the rate of annotated protein sequences in UniProt^1^ follows a linear trendline. This means that a gap is rapidly opening between raw sequence reads vs. annotated sequences (Figure 1). To meet this challenge, artificial intelligence (AI) algorithms have emerged as promising alternatives for the automated assignment of functions of uncharacterized proteins^3^. These models offer the promise for high quality automated functional annotation of sequenced genomes^3–5^. Recently developed methods such as CLEAN^5^, DeepECtransformer^6^ and ProteInfer^4^ have enabled accurate <u>E</u>nzyme <u>C</u>ommission (EC) number recapitulation by leveraging pretrained <u>p</u>rotein <u>L</u>anguage <u>M</u>odels^7,8^ (pLM) and deep learning algorithms. However, quantification of enzyme activity is still largely dependent on costly and time-consuming biochemical assays. Such approaches cannot keep up with the torrent of raw sequence reads leaving most computationally identified enzymes uncharacterized in terms of their kinetics despite significant progress in high throughput screening capacity ^9,10^. Therefore, predictive models that enable quantitative annotation of enzyme kinetics could be enabling for enzyme characterization in the same manner that recent fold prediction algorithms^7,11^ have become for structure prediction. Even approximate estimates of enzyme kinetics on a given substrate can be very important for a diversity of tasks ranging from starting point enzyme selection in directed evolution for protein engineering^12,13^, biosynthetic or biodegradation pathway pre-screening^14,15^, or initialization in the parameterization of kinetic models of metabolism^16^. Enzyme engineering efforts often rely on evolutionary methods such as directed evolution that aim to rachet up enzyme activity and/or selectivity. The selection process of the starting enzyme that undergoes directed evolution can be informed based on computationally derived enzyme kinetic estimates. *De novo* enzyme kinetic parameter prediction can also inform pathway assembly algorithms^17^ aimed at designing entire retro-biosynthetic routes for biochemical synthesis. Kinetic parameter predictions can be used to avoid alternatives with poor enzyme turnover or enzymes that exhibit strong product inhibition accelerating the discovery of more catalytically efficient routes. Finally, kinetic models, by relating enzyme kinetics to the concentration of metabolites and enzyme levels within a cell, can be used to both describe and redesign metabolism^18^.

**Figure 1.**
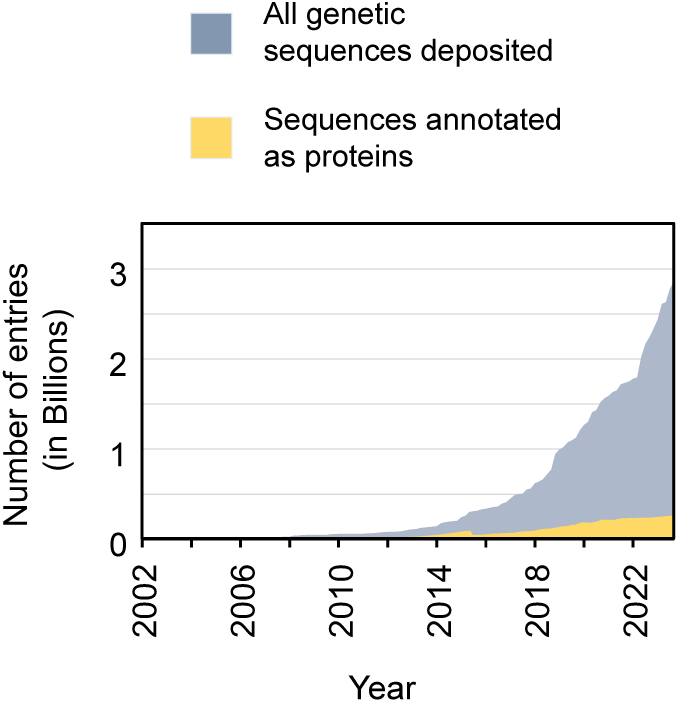
Growth of (a) genetic and (b) protein sequences over the past two decades as deposited in the World Genome Sequence (WSG) database and the UniProt database respectively.

Advances in automated functional annotation of proteins have enabled building metabolic models with a genome-wide coverage of cellular metabolism^19,20^. However, efficient kinetic parameterization to match observed fluxomic, proteomic and/or metabolomic datasets remains a bottleneck^21^. The use of reliable estimates for *in vitro* enzyme kinetic properties could accelerate convergence by serving as initializations of enzyme parameters^22^. These are but a handful out of the many applications that reliable enzyme parameter prediction could impact.

The catalytic turnover number and the Michaelis constant are key parameters of the Michaelis-Menten kinetics which is the universally accepted biochemical assay for quantitative assessment of enzyme function^23^. The turnover number, *k_cat_*, is the *speed* of an enzyme, the maximal number of molecules of substrates converted to products per active site per unit time. The Michaelis constant, *K_m_,* is equivalent to the concentration of a substrate at which the enzyme operates at half of its maximum catalytic rate qualitatively describing the binding affinity between the enzyme-substrate pair. Since enzymes have evolved to cater a wide array of cellular functions, they catalyze diverse chemical transformations and hence operate with a broad range of *k_cat_* and *K_m_* values^24^. In the presence of competitive or non-competitive inhibitors, the equivalent value of *K_m_* can be obtained using inhibition constants (*K_i_*). Databases such as BRENDA^25^ and SABIO-RK^26^ contain hundreds of thousands of *in vitro* kinetic measurements manually curated from primary research literature (Supplementary Table S1). Several previous studies have focused on developing ML models for *k_cat_* and *K_m_* prediction by using these database entries as training data^27–30^. Li. et. al^28^ developed *DLKcat*, by training a deep learning model on a dataset of 16,838 *k_cat_* values of both natural and engineered enzymes across various species. They used a convolutional neural network (CNN) architecture to extract features of enzyme-sequence motifs and a graph neural network (GNN) to extract substrate features using their 2-dimensional (2-D) connectivity graphs. Kroll. et. al. trained a gradient-boosted tree model, *TurNup*^27^, using language model features of enzymes’ amino acid sequences along with reaction fingerprints for *k_cat_* prediction using a dataset of 4,271 *k_cat_* measurements. Although *TurNup* was trained on much smaller dataset, they achieved a better generalizability compared to *DLKcat* on test enzyme sequences dissimilar to training sequences (out-of-distribution test examples)^27^. More recently, Yu et. al. developed *UniKP*^30^ for ML prediction of *k_cat_, K_m_* and *k_cat_/K_m_* values by training on previously curated datasets^28,29^. They trained a tree-ensemble regression model by utilizing pre-trained language models^8^ for extracting features of both enzymes and substrates. *UniKP* demonstrated an improved performance for *k_cat_* prediction compared to *DLKcat* on in-distribution tests, however, no out-of-distribution examples were tested. Currently, *TurNup* is the only prediction framework that is systematically evaluated on out-of-distribution tests for *k_cat_* prediction and outperforms *DLKcat* in this aspect presumably due to the use of pre-trained language model features.

Unlike *k_cat_* values that are not directly relatable to the physical properties of the substrate, *K_m_* values have been shown to be correlated with their molecular mass and hydrophobicity^31^. Kroll et. al.^29^ developed a *K_m_* prediction model using a gradient-boosted tree algorithm by training on 11,675 *in vitro* measurements of natural enzyme-metabolite pairs. They used a protein Language Model (pLM), UniRep^32,21^ for extracting numerical representations of the enzyme and a task specific graph neural network derived fingerprints combined with the molecular mass and hydrophobicity properties as features for metabolites. Yu et. al.^30^ also trained a *K_m_* prediction model within the UniKP framework using the same training dataset utilizing a more recently developed pLM, ProtT5^8^ for extracting enzyme sequence features. They demonstrated a similar performance as Kroll et.al^30^. Notably, both these existing models for *K_m_* prediction are only evaluated on in-distribution sequences (i.e., test enzyme sequences that are not explicitly excluded from those of training datasets). Relatively fewer ML models are available for *K_i_* prediction of enzyme-inhibitor pairs with most of them focused at predicting IC50 values of drug-target pairs^33,34^.

Existing studies for machine learning *in vitro k_cat_* and *K_m_* values either use BRENDA^25^, SABIO-RK^26^, UniProt^1^ or a combination of these to curate their training datasets from known measurements of kinetic parameters. However, there is a lack of complete annotations in the databases for all entries leaving significant gaps in the amount of learnable data. For example, even though there exist about 87k, 176k and 46k entries for *k_cat_*, *K_m_* and *K_i_* measurements, respectively in BRENDA (Release 2022_2), many are not annotated with the corresponding enzyme sequences and/or substrate information. Owing to this, training datasets used by existing works vary significantly depending on how they handle entries with missing information. This has prompted most studies to use small, filtered subsets of the available data to mitigate this effect. For example, *TurNup* for *k_cat_* prediction is trained only on 1,192 enzyme types (unique EC numbers) while the current biochemical databases contain *k_cat_* values for over 3,000 enzyme types (Supplementary Table S2). Many studies have also imposed arbitrary exclusion criteria with the goal of reducing the effect of noisy measurements^27,29^. While such filtering may in part reduce the effect of noise, it could also potentially lead to information loss, biasing, and overfitting to the training datasets especially when high-dimensional deep learning architectures are used. Filtered-out entries often correspond to infrequently occurring metabolite entries. Since they correspond to a large fraction (i.e., up to ∼ 40-70%, Supplementary Table S3) of available data entries, their omission can become a missed opportunity for ML algorithms to learn on rarely seen data and expand coverage of generalizable latent spaces. Another notable source of incongruency between different datasets is the mapping process adopted of substrate names to their respective chemical connectivity information using SMILES^35^ strings. Existing studies use either of, or a combination of PubChem^36^, KEGG^37^ or ChEBI^38^ databases to map substrate names to the respective database identifiers and subsequently retrieve SMILES strings leading to divergent results in some cases thus precluding a fair comparison across machine learning frameworks. This motivates the need for both systematic data curation pipelines and standardized training datasets with expanded enzyme and substrate scope.

ML models trained on noisy datasets can lead to potentially unreliable predictions especially when challenged with inputs significantly different from those that the model is trained on. Predictive models that display good performance on enzyme sequences that are under-represented in training datasets require that the models have learnt generalizable information encoded in latent spaces instead of overfitting to nuances/noise present in the training data.

Existing ML models for *k_cat_* or *K_m_* prediction use traditional regression approaches by minimizing the mean-squared-error between training data and thus output deterministic (single valued) enzyme parameter predictions. These predictions lack any confidence metric information. In contrast, probabilistic regression approaches can output predictions as gaussian distributions (including a mean and a variance) which has the potential to offer guardrails on the reliability of predictions. Such methods have been recently explored in the molecular property prediction domain where similar challenges with datasets exist^39^.

Here we introduce the comprehensive ML framework, CatPred, for enzyme kinetic parameter prediction that addresses many of the aforementioned challenges. We first assembled an expanded set of benchmark datasets, CatPred-DB, for training and evaluating ML models using *in vitro* kinetic measurements of *k_cat_, K_m_* and *K_i_* extracted from both BRENDA and SABIO-RK databases. Using these datasets, we train deep learning models utilizing features of different levels of complexity – enzyme sequence level (using sequence-attention and pLM features), and enzyme structure level equivariant graph neural network (E-GNN)^40^ derived features. Substrate representation by CatPred relies on a graph neural network approach previously shown to be promising for a wide range of molecular property prediction tasks^41^. By leveraging a probabilistic regression approach^39^ that simultaneously learns to output means and variances of predictions, CatPred provides confidence estimates to its predictions. We systematically evaluated the predictive performances of CatPred on test datasets containing both in-distribution and out-of-distribution enzyme sequences (different from sequences encountered during training). Our results show that pLM derived features are necessary for achieving good predictive performances on out-of-distribution enzymes. CatPred performs favorably in a range of benchmarks compared to existing approaches while also offering uncertainty quantifications to its predictions.

## Results

### Generation of benchmark datasets CatPred-DB of *in vitro* enzyme kinetic parameters

CatPred-DB consists of a set of comprehensive benchmark datasets for training ML models, one each for *k_cat_*, *K_m_* and *K_i_ in vitro* measurements. We used data from the BRENDA release 2022_2 and data from the SABIO-RK as of November 2023. Initially, we parse the databases to identify entries containing essential information, including at least one kinetic parameter value (*k_cat_*, *K_m_*, or *K_i_*), the enzyme type (EC number), the organism of enzyme’s origin, and the names of reactants and products. To maintain the accuracy of organisms’ names, we retain entries only if they are listed in the NCBI Taxonomy database^42^. We then mapped each entry to the enzyme’s amino acid sequence identifier using the UniProt database (Methods for details). We excluded entries that lack one or more of these annotations or if any of these annotations are incomplete. Finally, each substrate name is used to obtain a canonical SMILES string that corresponds to the 2D atom connectivity. If there exist multiple measurements of any parameter belonging to an enzyme-sequence and substrate-SMILES pair, then the maximum (for *k_cat_*) and the geometric mean (for *K_m_* and *K_i_*) value, respectively is retained. The selection of the maximum value for *k_cat_* value is carried out because it likely maps to the optimal growth conditions (i.e., temperature, pH, etc.). In contrast, *K_m_* and *K_i_* values are more directly associated with the enzyme-substrate/inhibitor affinities rather than on the experimental conditions. The use of the geometric average implies an arithmetic averaging of the logarithmically transformed values used in the training process. The selection of a unique value for the enzymatic parameters is needed to safeguard against the ML method attempting to learn significantly different outputs for the same inputs which can result in instabilities during training.

CatPred-DB contains 23,197 *k_cat_*, 41,174 *K_m_* and 11,929 *K_i_* measurements spanning thousands of unique enzymes, organisms, and substrates (Table 1). Each entry in CatPred-DB is also mapped to a predicted 3D-structure of the corresponding enzyme using AlphaFold-2.0 database^11^. In the absence of a 3D structure in the AlphaFold database, we used ESMFold^7^ to carry out structure prediction. The coverage statistics of CatPred-DB contrasted with other efforts^28–30^ are summarized in Table 1. Notably, CatPred-DB has a significantly expanded enzyme sequence space (up to 60% new sequences introduced) in comparison to the existing ML datasets for *k_cat_* and *K_m_*. New sequences span widely across enzyme classes with no biases for specific EC classes (Figure 2b). Moreover, *k_cat_* and *K_m_* entries in CatPred-DB have broader coverages compared to existing ML datasets across all the enzyme families as per the EC level 1 (Figure 2c). Therefore, we envision that the enhanced sequence and EC classification coverage would make CatPred-DB a useful resource to the community for aiding systematic development and benchmarking of ML models for enzyme kinetic parameter prediction.

**Figure 2.**
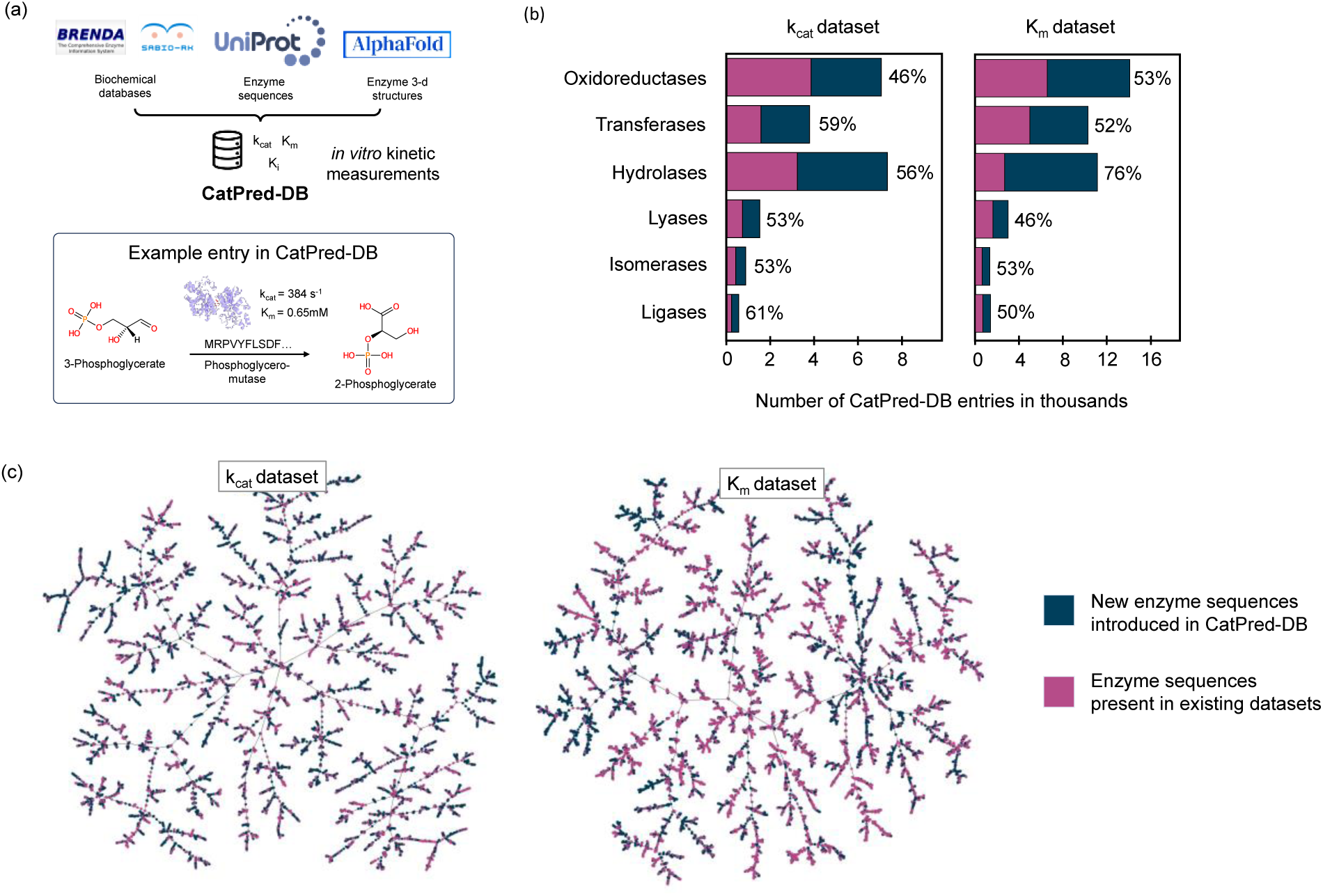
**(a)** CatPred-DB is a comprehensive collection of benchmark datasets for *k_cat_*, *K_m_* and *K_i_* including *in vitro* measurements of enzymatic reactions curated from BRENDA and SABIO-RK databases. For each enzymatic reaction, the datasets contain complete annotations of the molecules involved in the reaction, the enzyme sequence, the AlphaFold2.0/ESMFold predicted enzyme structure and the associated kinetic parameters. (b) Bar plot of the number of entries in the CatPred-DB - *k_cat_* and *K_m_* datasets grouped by their Enzyme Classification (EC level 1). Each bar is divided into two differently colored portions corresponding to enzyme sequences newly introduced in CatPred-DB (blue) and to enzyme sequences present in existing datasets (magenta). The percent entries on top of each bar show the newly added sequences. (c) The enzyme sequence latent space plots of CatPred-DB’s *k_cat_* and *K_m_* datasets visualized using the ESM-2 protein Language Model (pLM) embeddings. The sequence embeddings are converted to k-nearest neighbor graphs (k=10) and visualized using the TMAP^56^ and Faerun^57^ libraries. Each point in the latent space plots corresponds to a single enzyme sequence and is colored according to whether it has been newly introduced in CatPred-DB (blue) or is present in existing datasets (magenta).

**Table 1.** Coverage statistics of CatPred-DB vs. other datasets of in vitro enzyme kinetic parameter measurements.

| Dataset | CatPred-DB |  |  | Existing datasets |  |
| --- | --- | --- | --- | --- | --- |
| | $k_{cat}$ | $K_m$ | $K_i$ | $k_{cat}$ (Li. et. al.<sup>28</sup>) | $K_m$ (Kroll et. al.<sup>29</sup>) |
| **Entries** | 23,197 | 41,174 | 11,929 | 17,010 | 11,722 |
| **Unique organisms** | 1,685 | 2,419 | 652 | 849 | N/A |
| **Unique Enzyme Classes (EC)** | 2,657 | 3,550 | 1,306 | 1,692 | 3,690<sup>#</sup> |
| **Unique enzyme sequences** | 7,183 | 12,355 | 2,829 | 3,219 | 6,990 |
| **Unique substrates** | 12,290 | 10,535 | 7,146 | 2,696 | 1,566 |

<sup>#</sup> Predicted Enzyme Classification (EC) numbers using CLEAN

### Overview of CatPred training framework

CatPred relies on the enzyme sequences/3D-structures along with the SMILES string of the corresponding substrates (reactants) as inputs and outputs machine-learned *in vitro* kinetic parameters. We used a concatenated SMILES string of all the reactant molecules for *k_cat_* prediction. For *K_m_* or *K_i_* prediction, the SMILES string corresponding to the relevant substrate is used. During training, the two sets of inputs are first transformed into their respective feature spaces through separate feature learning modules (Figure 3a). For enzyme feature learning, *CatPred* makes use of three approaches that successively add to the detail of description: (1) <u>Seq</u>uence <u>Att</u>ention (Seq-Att) (2) <u>p</u>rotein <u>L</u>anguage <u>M</u>odel (pLM) features, and finally (3) 3D-structure features (Figure 3c). This is carried out to properly delineate the respective contribution to improved prediction of more sophisticated encodings. For substrate feature learning, CatPred utilizes the extensively benchmarked <u>D</u>irected <u>M</u>essage <u>P</u>assing <u>N</u>eural <u>N</u>etworks^41^ (D-MPNN). D-MPNNs transform SMILES strings to 2D-graphs of atoms with bond connectivity and learn their aggregated representations using graph convolution operations^41^ (Figure 3b). For the derivation of sequence attention (Seq-Attn) features, the amino-acid sequences of enzymes are encoded into numerical representations using the rotary positional embeddings^43^ akin to the encoding layer used for training the ESM-2 pLM^7^. The encoded numerical representations are then transformed using self-attention layers^44^ to capture dependencies and relationships across the length of enzyme sequences (Figure 3a). The pLM features are extracted by using the ESM-2^7^ (<u>E</u>volutionary <u>S</u>cale <u>M</u>odeling) model pretrained on the Uniref50 dataset. The 3D structural features are extracted using the <u>E</u>quivariant <u>G</u>raph <u>N</u>eural <u>N</u>etworks (E-GNN^40^) that operate on amino acid residue graphs. We integrated E-GNN from Greener et. al.^45^ that has been pre-trained using a supervised contrastive learning for embedding protein structures into a low-dimensional latent space (Figure 3a). The pre-trained E-GNN’s latent space clusters the embeddings of similar protein structures together whereas separating dissimilar ones away from one another ^45^. We reasoned /that using these E-GNN derived embeddings as features within CatPred can complement the sequence-attention and pLM features. Enzyme features learnt through these modules (Seq-Attn, pLM, E-GNN) are concatenated along with the substrate features from D-MPNNs and used to predict the respective targets (log10-transformed kinetic parameters). CatPred uses a probabilistic regression approach^46^ and therefore provides kinetic parameter predictions as distributions characterized by both a mean and a standard deviation, rather than single value predictions. Specifically, the concatenated enzyme and substrate features are fed into a fully connected neural network which outputs a mean and variance for each input (Figure 3c). The network is trained using a negative log likelihood (NLL) loss function with respect to the CatPred-DB’s

**Figure 3.**
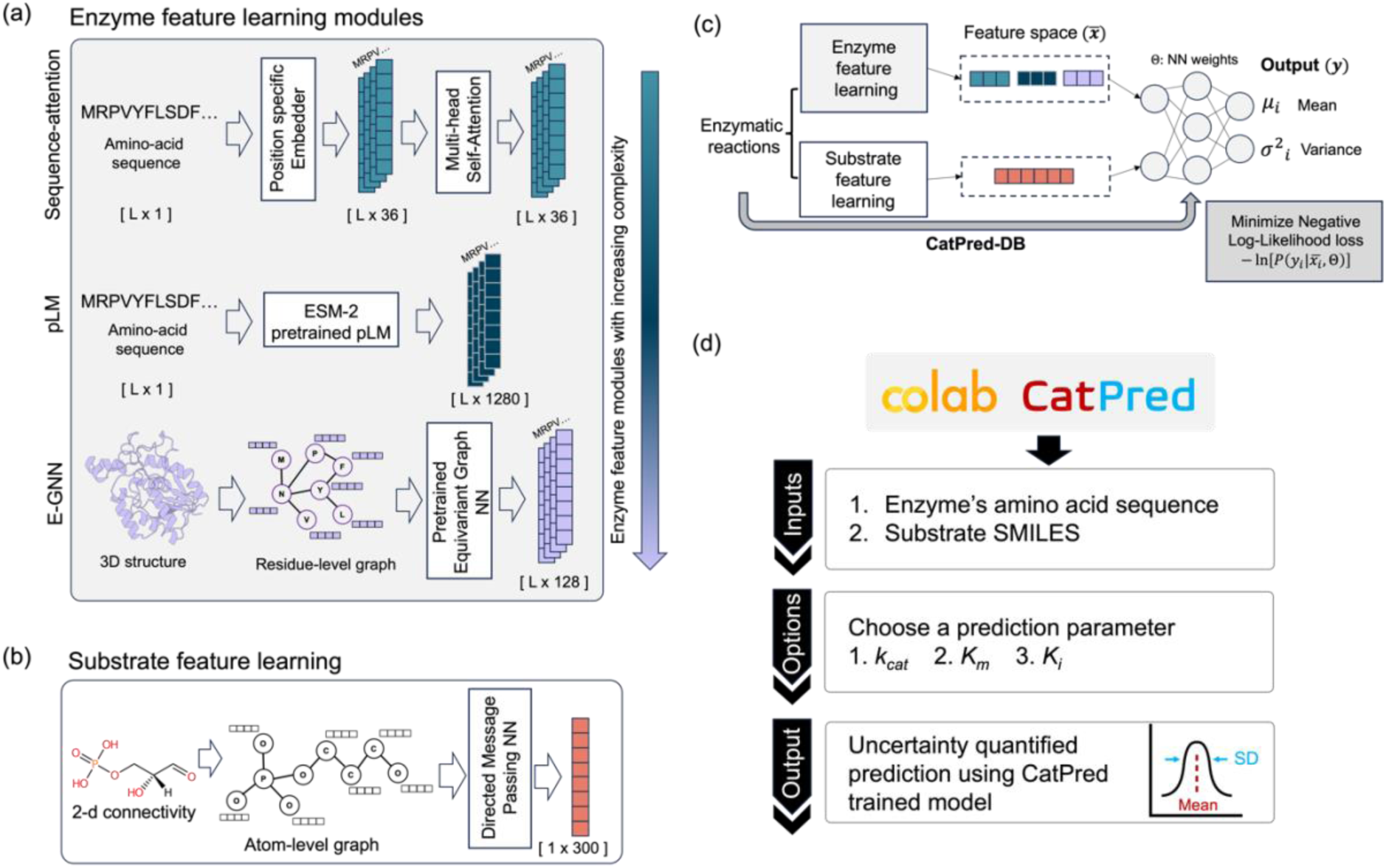
CatPred framework for training probabilistic regression models for enzyme kinetic parameter prediction using substrate and enzyme features. **(a)** Enzyme feature learning is carried out using three different modalities with increasing level of detail. The <u>Seq</u>uence-<u>Att</u>ention (Seq-Att) module learns features of amino-acid embeddings using multi-head attention layers. The pLM module uses features extracted from a pre-trained <u>p</u>rotein <u>L</u>anguage <u>M</u>odel (pLM). The <u>E</u>quivariant <u>G</u>raph <u>N</u>eural <u>N</u>etwork (E-GNN) module extracts features of 3d structures of enzymes by employing equivariant graph neural networks on their amino-acid level graphs. **(b)** Substrate feature learning is carried out using <u>D</u>irected <u>M</u>essage <u>P</u>assing <u>N</u>eural <u>N</u>etworks (D-MPNN) that extract molecular representations by leveraging 2D atom-bond connectivity graphs. **(c)** CatPred models are trained on CatPred-DB datasets utilizing both substrate and enzyme feature learning modules with a probabilistic regression approach. The enzyme and substrate features are input to a fully connected neural network that predicts the kinetic parameters as outputs in the form of Gaussian distributions characterized by their respective means (*μ*) and variances (*σ*^2^). **(d)** CatPred production models are made available through the Google-Colab interface for ease of access. The inputs are the substrate SMILES and either enzyme sequence or structure along with a choice of kinetic parameter for prediction. The interface then loads the respective trained models and outputs uncertainty quantified kinetic parameters in terms of a predicted mean and standard deviation (SD).

For each dataset in CatPred-DB, the CatPred framework is used to train ML models that minimize a negative log-likelihood loss^46^ (Methods for details) of the predicted distributions to the corresponding target values. Each CatPred-DB dataset is randomly split into 80-10-10 proportions for training-validation-testing, respectively. Because CatPred involves using both enzyme sequences/structures and substrate SMILES as inputs, the splitting is carried out so as no enzyme-substrate pair is repeated across different partitions. Adjustable hyper-parameters in the framework are either fixed to default values or optimized by evaluating trained CatPred models on the validation sets (Methods). The optimized hyperparameters are used to train the final models CatPred-*k_cat_*, CatPred-*K_m_* and CatPred-*K_i_* using the training and validation sets and evaluated on the testing sets (see below). Production models trained on the full datasets are made available for easy access through the Google Colab interface which can be used without the requiring any local installation or specialized hardware (Figure 3d).

### Evaluation of trained CatPred models

Trained CatPred models were evaluated on two test sets – (1) “held-out” test set and (2) “out-of-distribution” test set. The evaluation criterion is based on the coefficient of regression (R^2^) which quantifies the fraction of data variance in the regression target that is captured by the predicted values. For each kinetic parameter, the held-out test sets are constructed to be randomly selected 10% in size subsets of the complete CatPred-DB dataset. As implied by their definition, the held-out test sets do not contain any enzyme-substrate pairs used for training the models. The out-of-distribution test sets are further subsets of the held-out test sets (approximately 12 to 15% thereof) with not only specific enzyme-substrate but all enzyme sequences (nearly) identical excluded from the training set (Figure 4a). By construction, any enzyme sequence in the out-of-distribution set is at most 99% identical (Methods) to any sequence in the training set. Therefore, prediction metrics achieved on the held-out test sets reflect the prediction fidelity for unseen enzyme-substrate pairs. Out-of-distribution test sets provide a more stringent prediction challenge by assessing prediction performance on unseen enzymes (even excluding enzymes within 99% in sequence identity).

**Figure 4.**
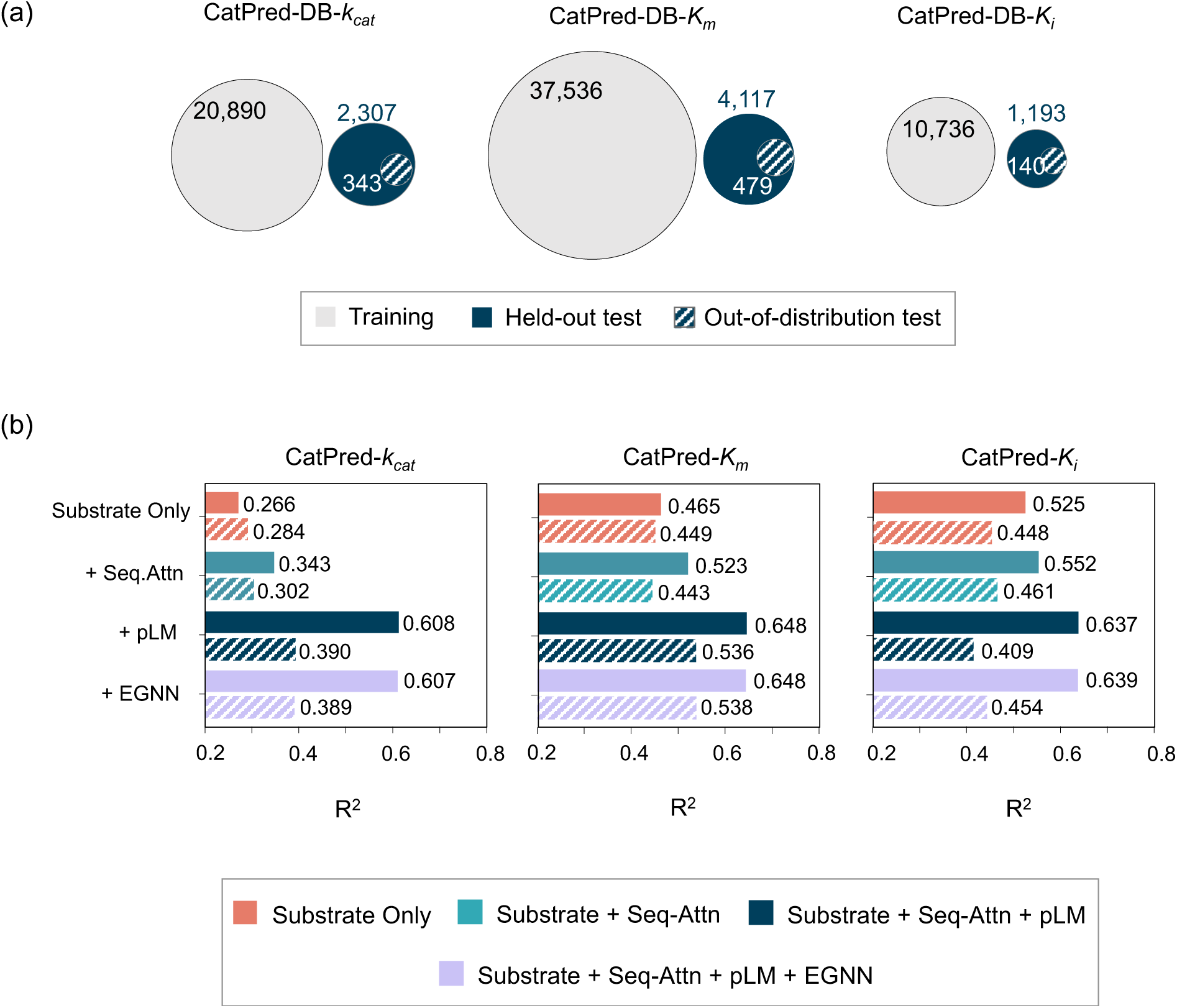
(a) CatPred-DB dataset sizes used for training, held-out test and out-of-distribution test are shown as Venn diagrams. (b) Coefficient of determination (R^2^) values obtained by trained CatPred models for kcat, Km and Ki prediction on held-out and out-of-distribution test sets. (a) by the models on (hold out) test sets (solid bars) and on (out-of-distribution) samples (patterned bars) are shown. The out-of-distribution samples are subsets of the full test-sets extrscted so as no enzyme sequence in the subset is more than 99% similar to any training sequence. ‘Substrate Only’ refers to CatPred models trained using only the substrate features; ‘Substrate+Seq-Attn’ (<u>Seq</u>uence <u>Att</u>ention) refers to CatPred models trained using substrate features and the Seq-Attn features; ‘Substrate+Seq-Attn+<u>pLM</u>’ (<u>p</u>rotein <u>L</u>angugae <u>M</u>odel) refers to CatPred models trained using substrate features along with both the Seq-Attn and pLM features; ‘Substrate+Seq-Attn+pLM+<u>EGNN</u>’ (<u>E</u>quivariant <u>G</u>raph <u>N</u>eural <u>N</u>etworks) refers to CatPred models trained using substrate features along with Seq-Attn+pLM and EGNN features.

We find that CatPred models that use substrate features along with both Seq-Attn and pLM features have the best performance across all three enzymatic parameters (Figure 4b). Notably, using only the substrate features leads to a reasonable performance for both *K_m_* and *K_i_* prediction (R^2^ of 0.465 and 0.525) at par with previous studies^29^. Even though inclusion of Seq-Attn features alone only slightly improves prediction performance, the combined addition of both Seq-Attn and pLM features leads to best “in-class” performance for *k_cat_*, *K_m_* and *K_i_* prediction with R^2^ values of 0.607, 0.648 and 0.637, respectively (Figure 4b). These metrics are at least as good or better than all existing ML models for predicting *k_cat_*^27,28,30^ and *K_m_*^29,30^ values respectively. It is worth noting that CatPred models that use 3D-structural features extracted from the E-GNN in addition to Seq-Attn and pLM features do not improve the prediction performance compared to only using Seq-Attn and pLM. The achieved R^2^ values were 0.607, 0.648 and 0.639 on the held-out test sets respectively for *k_cat_*, *K_m_* and *K_i_* (Figure 4b).

Importantly, CatPred models retained strong prediction performance even on “out-of-distribution” test sets for *K_m_* (R^2^ = 0.536) and somewhat less accurate for *k_cat_* and *K_i_* (R^2^ = 0.390 and 0.409 respectively) (Figure 4b). We observe that while adding Seq-Attn features leads to improved performance for *k_cat_* and *K_m_* predictions, the improvements are not as pronounced on out-of-distribution sets. This suggests that even though the self-attention layers in Seq-Attn can successfully encode enzyme sequences by extracting local and global patterns, they cannot account for higher-order relationships across sequences that are necessary for generalization to unseen protein sequences. ESM-2 pLM can capture such features and have already proven capable of encoding evolutionarily rich semantics of protein sequences^7,47^ explaining their good performance on out-of-distribution samples.

We found that adding Seq-Attn+pLM features leads to a reduction in the R^2^ value for *K_i_* prediction on out-of-distribution test sets when compared to adding only Seq-Attn features. This seemingly surprising finding is likely due to overfitting on the relatively small *K_i_* dataset (approximately four-fold smaller than *K_m_* dataset, see Table 1) using high dimensional pLM features. This calls for an expansion to the size of the *K_i_* dataset in the future. It is worth noting that CatPred performs (R^2^ = 0.39) comparably with TurNup (R^2^ = 0.40) on out-of-distribution samples for *k_cat_* prediction. To the best of our knowledge, CatPred is the only available predictive model for *K_m_* and *K_i_* prediction that is evaluated on out-of-distribution samples.

Recently, Kroll et al.^27^ reported that the DLKcat model for *k_cat_* prediction showed a diminishing performance as a function of the similarity of test enzyme sequences to those of the training set indicating that the DLKcat model might have “memorized” the training dataset instead of “learning” meaningful patterns. They showed that the DLKcat model exhibited poor predictive performance (R^2^ = −0.61) on sequences that are significantly dissimilar compared to those in the training set. Motivated by the need to avoid such a prediction behavior, we systematically assessed the reduction in prediction performance of CatPred models as the test sets become more and more dissimilar to the training set. This analysis revealed that CatPred models for *K_m_* prediction maintain robust performance with an R^2^ value of 0.48 even on out-of-distribution test sets with sequence similarities less than 40% when pLM features are enabled (Figure 5b). Prediction by CatPred for *k_cat_* values remain reasonable (i.e., R^2^ = 0.33) even down to a seq. id. cutoff of 40% (Figure 5a) with the contribution of pLM encodings being even more pronounced. This suggests that the CatPred models for *k_cat_* and *K_m_* (with pLM features) have learnt generalizable enzyme attributes that go beyond sequence similarities. In contrast, for CatPred-*K_i_* the benefit of using pLM features is not realized presumably due to overfitting caused by the relatively small training set size. However, using only Substrate and Seq-Attn features, a good predictive performance is reached for *K_i_* with an R^2^ value of 0.42 even on the test set with <40% similarity to training sequences (Figure 5c). Also, for CatPred models using E-GNN features, the corresponding R^2^ values on the out-of-distribution test sets were 0.389, 0.538 and 0.454 for *k_cat_*, *K_m_* and *K_i_* respectively (Figure 4b) indicating no significant improvement over using only Seq-Attn+pLM features. Therefore, the production CatPred models accessible through our Google Colab interface (Figure 3d) are based on Substrate+Seq-Attn+pLM for *k_cat_* and *K_m_* and only Substrate+Seq-Attn for *K_i_*. Also, all further mentions of CatPred-models throughout the manuscript refer to these models unless otherwise explicitly specified.

**Figure 5.**
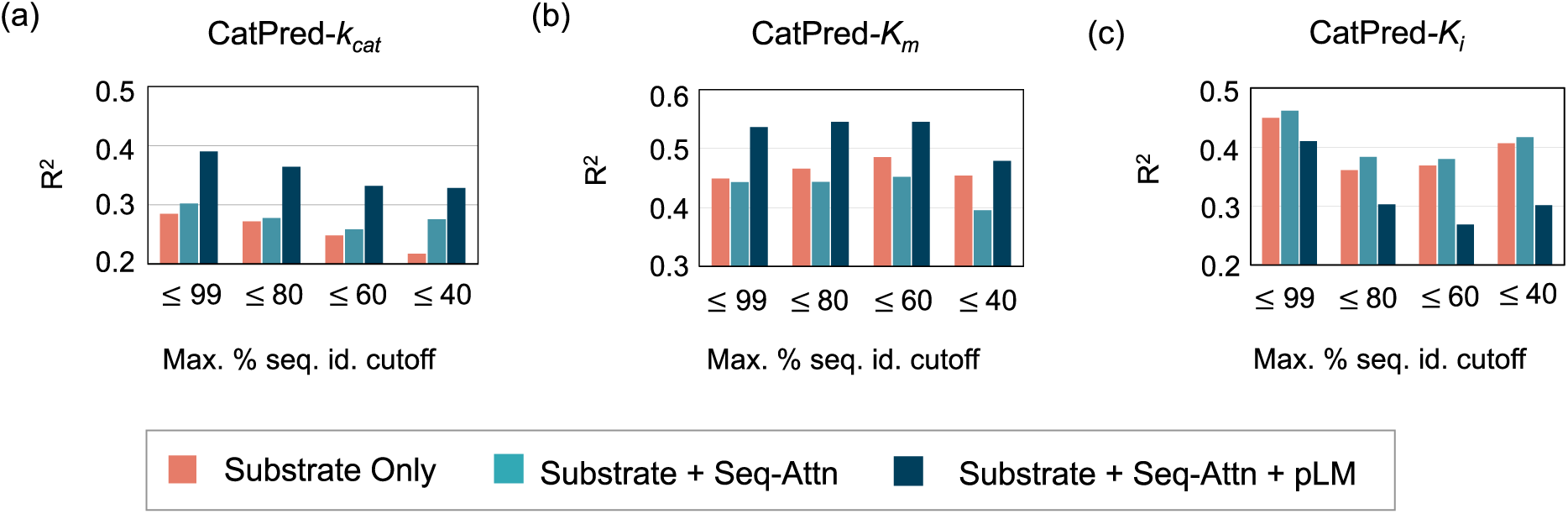
Evaluation of trained CatPred models on out-of-distribution sets with decreasing enzyme sequence similarities to training sequences for (a) *k_cat_* (b) *K_m_* and (c) *K_i_* respectively. Each group on X-axis indicates the coefficient of determination (R^2^) obtained on subsets of held-out tests selected using a maximum percent sequence identity cutoff (Max. % seq. id. cutoff) to training sequences. ‘Substrate Only’ refers to CatPred models trained using only the substrate features. ‘Substrate+Seq-Attn’ (<u>Seq</u>uence <u>Att</u>ention) refers to CatPred models trained using substrate features and the Seq-Attn features. ‘Substrate+Seq-Attn+pLM’ (<u>p</u>rotein <u>L</u>anguage <u>M</u>odel) refers to CatPred models trained using substrate features along with both the Seq-Attn and pLM features.

In the analyses described above we used R^2^ as the sole metric of prediction quality. We have repeated almost all assessments and Figures using the mean absolute error (MAE) metric (Supplementary Figure S1) obtaining the same trendlines. However, neither R^2^ nor MAE provide immediate feedback to the user as to whether the predicted value for the enzyme parameter is likely to be “order of magnitude” accurate or not. Motivated by the need to provide such a metric, we introduced a new metric termed p_1mag_ defined as the percent of test predictions that are within one order (+/-) of magnitude error. We choose the relatively large window of acceptance of one order of magnitude as enzyme kinetic parameters span multiple orders of magnitude. Table 2 shows the performance evaluation of CatPred models in terms of R^2^, MAE and p_1mag_. Results indicate that approximately 80%, 87% and 70% of held-out test predictions fall within an order of magnitude error for *k_cat_*, *K_m_* and *K_i_* predictions, respectively. They drop to 63.5%, 82.7% and 58.6% when evaluated on the out-of-distribution test sets. The p_1mag_ metric provides a confidence level metric evaluated for an entire subset of measurements. We next describe how one could directly use the variances predicted by the probabilistic regression model in CatPred to infer confidence values for each prediction separately. Reliable confidence estimates can help segregate predictions with small errors from those with larger ones.

**Table 2.**
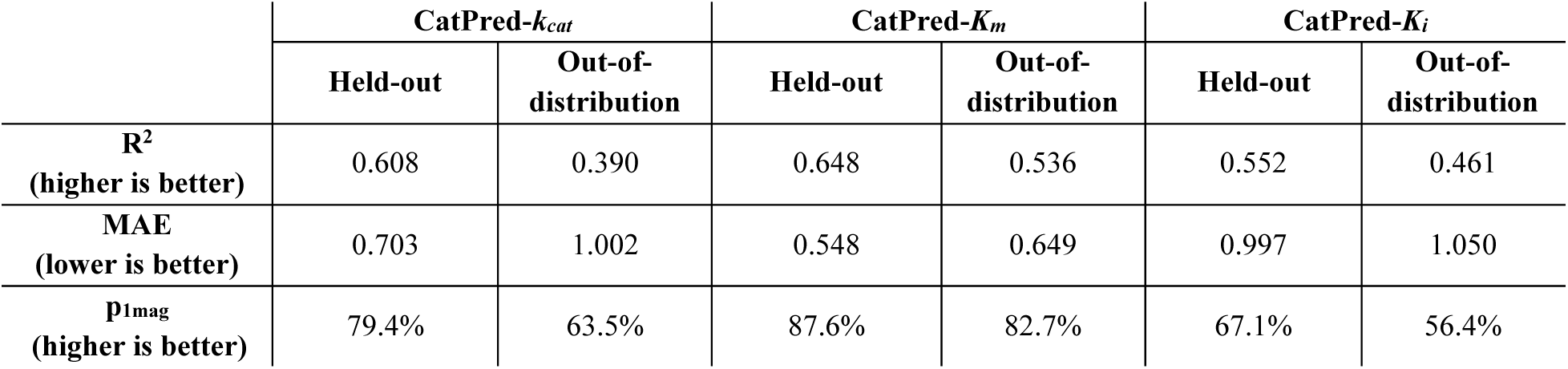
The performance metrics obtained by CatPred models as quantified using the coefficient of regression (R^2^), the mean absolute error (MAE), and the percent of predictions within test sets that are within one order of mangintude error (p_1mag_). Prediction metrics obtained on both held-out test sets and out-of-distribution sets are listed.

| | CatPred-$k_{cat}$ | | CatPred-$K_m$ | | CatPred-$K_i$ | |
| --- | --- | --- | --- | --- | --- | --- |
|  | Held-out | Out-of-distribution | Held-out | Out-of-distribution | Held-out | Out-of-distribution |
| **$R^2$**<br>(higher is better) | 0.608 | 0.390 | 0.648 | 0.536 | 0.552 | 0.461 |
| **MAE**<br>(lower is better) | 0.703 | 1.002 | 0.548 | 0.649 | 0.997 | 1.050 |
| **$p_{1mag}$**<br>(higher is better) | 79.4% | 63.5% | 87.6% | 82.7% | 67.1% | 56.4% |

### Uncertainty estimates for predictions using CatPred models

Regression models described in the earlier section for training ML models of *k_cat_* and *K_m_* relied on a mean-squared error loss function^27–30^. This approach precludes quantifying the level of uncertainty of predictions for individual enzyme-substrate pairs. The metrics such as R^2^, MAE or f_1mag_ are assessed for the entire evaluation set (i.e., held-out or out-of-distribution) and not for individual predictions. Either lack of measurements or noisy data can adversely affect predictions for enzyme-substrate pairs. This implies that not all predictions would have the same fidelity.

Using a probabilistic description allows CatPred to quantify the uncertainty in prediction for individual enzyme-substrate pairs. There are two sources of encountered uncertainty (i.e., aleatoric and epistemic^39^). Aleatoric uncertainty arises from noise in the training data due to randomly occurring experimental error. This leads to uncharacteristic fluctuations in the value of the output even for small changes in the input (Figure 6c). Epistemic uncertainty arises due to the lack (or insufficiency) of training data in certain regions of the input space (Figure 6c). Aleatoric uncertainty can be captured using the probabilistic regression approach used in CatPred (Methods for details). By training the neural networks using a negative log likelihood (NLL) loss function, each CatPred model estimate is a Gaussian distribution characterized by a mean and a variance (Figure 6a). Epistemic uncertainty on the other hand, requires estimating the variance in prediction from an ensemble of identical neural network models trained using different initializations (Figure 6b). Individual models in the ensemble would provide dissonant predictions for inputs corresponding to regions with insufficient training data (Figure 6c). The extent of the disagreement thus quantifies the associated epistemic uncertainty. For each kinetic parameter prediction made by CatPred, the combined uncertainty (sum of aleatoric and epistemic contributions) is provided (Figure 6b). The aleatoric uncertainty is quantified as the square root of the arithmetic mean of ensemble variances (Figure 6b) whereas the epistemic uncertainty is the sample standard deviation of the ensemble means (Figure 6b, also see Methods). It is important to note that because the model training is performed using log10-transformed kinetic parameter values, the corresponding standard deviations estimated are also on a log10-scale (Methods for details). A similar uncertainty description framework was used before in molecular property prediction^39^.

**Figure 6.**
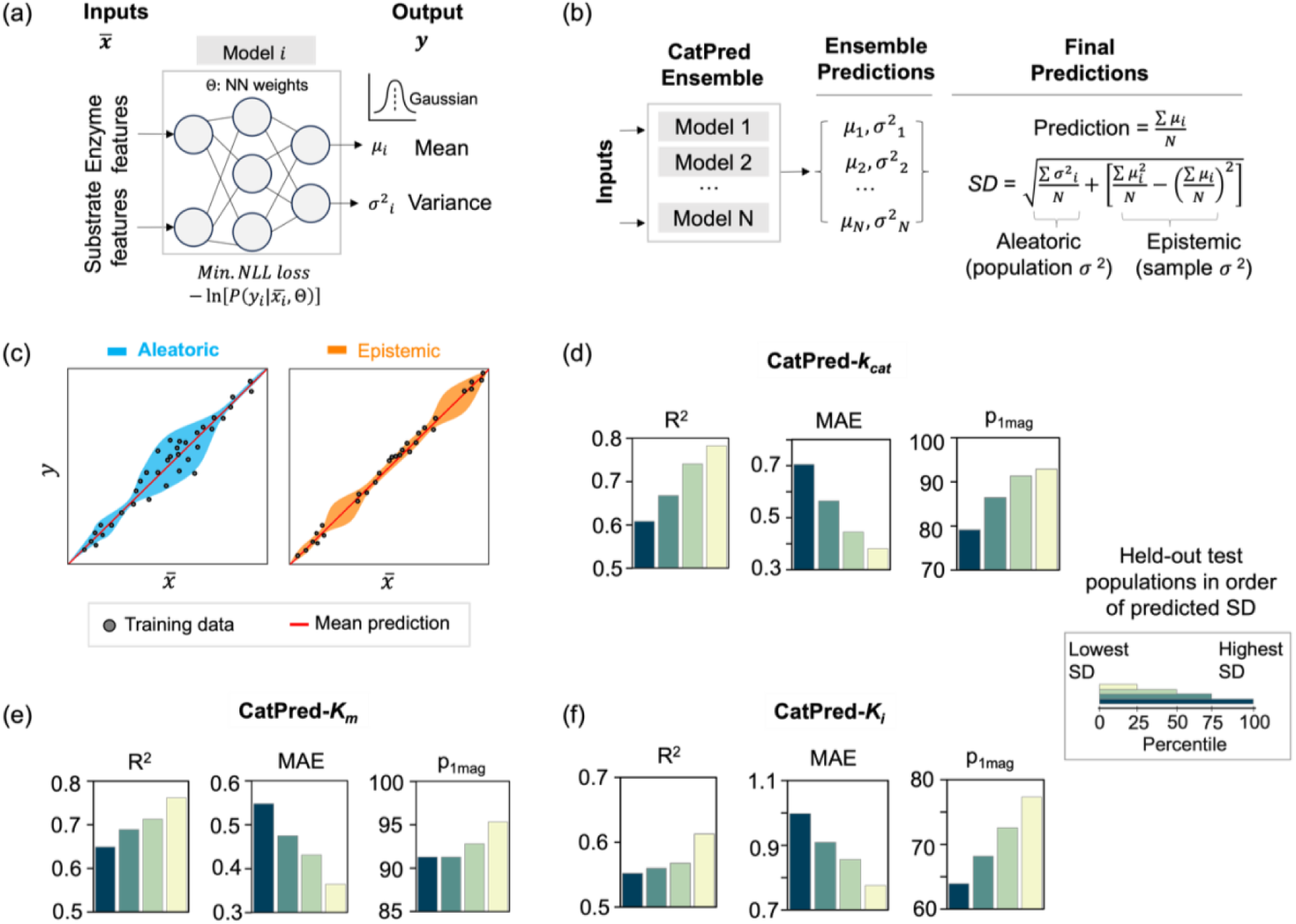
(a) CatPred uses as inputs enzyme and substrate features and outputs kinetic parameters as Gaussians distributions characterized by a mean and a variance. When training an ensemble of models, ‘Model *i’* corresponds to the *i*^th^ set of randomly initialized weights. (b) Uncertainty prediction pipeline in CatPred. An ensemble of N independent models (each with a unique set of randomly initialized weights) is trained for each prediction target *k_cat_*, *K_m_* and *K_i_*. Each model outputs a mean and a variance for a given set of inputs. The final prediction is the arithmetic average of ensemble means and the final uncertainty is the sum of aleatoric and epistemic contributions. (c) Schematic depicting the two kinds of uncertainties: aleatoric and epistemic. Aleatoric uncertainty is higher in areas with larger spread of the regression target variable, *y*, with respect to the input latent space *x*ҧ. Epistemic uncertainty is higher in areas with absence of knowledge of *y* within the training data. The circles in plots refer to training data and the red line denotes the mean prediction by trained models. The performance metrics achieved by (d) CatPred-kcat, (e) CatPred-Km and (f) CatPred-Ki models on sub populations of the held-out tests binned in order of their predicted uncertainty values (sum of aleatoric and epistemic uncertainty). Each colored bar denotes a sub population of the held-out set with uncertainty less than the 100^th^ (Blue), 75^th^ (Dark Green), 50^th^ (Light Green), and 25^th^ (Light yellow) percentile respectively. Within each figure, the subplots show the obtained co-efficient of regression (R^2^), mean absolute error (MAE) and percent of predictions within one oder of magnitude error (p_1mag_).

We first verified if the predicted uncertainty values are consistent with the absolute errors for predictions made by the CatPred trained models on held-out test sets. The goal was to ensure that the predicted uncertainties can be used to discriminate between high from low confidence predictions. To this end, the held-out test sets were partitioned in four subsets each consisting of predictions with uncertainty values less than the 100^th^, 75^th^, 50^th^ and 25^th^ percentile, respectively. This means that each subset becomes progressively enriched with predictions of higher confidence. Performance metrics R^2^, MAE and p_1mag_ are calculated separately within each subset (Figure 6 (d) –(f)). We perform these analyses on CatPred production models i.e., based on Substrate+Seq-Attn+pLM for *k_cat_* and *K_m_* and only Substrate+Seq-Attn for *K_i_*. We observe that the prediction metrics monotonically improved when held-out subsets with smaller predicted uncertainties are assessed (Figure 6 (d) –(f)). We note that R^2^ values for the (25^th^ percentile) set are improved to 0.78, 0.76 and 0.61 for CatPred-*k_cat_*, CatPred-*K_m_* and CatPred-*K_i_* models, respectively. Similarly, the MAE drops by approximately ∼36% for the 25^th^ percentile set compared to the 100^th^ percentile set. This trend is also reflected by the increase in p_1mag_ values (Figure 6 (d) – (f)) showing that more than 90% of predictions in the highest confident subset (i.e., 25^th^ percentile subset) are within an order of magnitude error for *k_cat_* and *K_m_* prediction. We also carried out this analysis for the out-of-distribution tests and we observed similar trends (Supplementary Figure S2). These results imply that the probabilistic description of CatPred correctly assigns lower standard deviations for predictions associated with higher confidence evaluation sets.

### Google Colab web interface for using CatPred

We developed an easy-to-use interface on Google Colab (https://tiny.cc/catpred) for accessing CatPred. This interface allows for remote computations in a web browser without requiring any local installation. The input to CatPred is the amino-acid sequence of the enzyme and the substrate SMILES string. In the case of *k_cat_* prediction, the substrate SMILES string must contain the concatenation of the SMILES strings associated with all reactants. As discussed previously this is needed as we discovered that not only the primary substrate but also the co-substrates (such as secondary substrates, cofactors etc.) contain information relevant to *k_cat_* prediction. Unsurprisingly, this is not the case for *K_m_* and *K_i_* where only substrate connectivity information is needed. Once the enzyme parameter of interest is chosen and the inputs are entered, they are validated for correct formatting. If the enzyme sequence contains characters other than the natural amino-acid alphabet or if the SMILES string is invalid, then an error prompt is displayed asking for re-entry of inputs. Once the inputs are validated, the relevant enzyme parameter prediction value along with the estimated uncertainty (contributions from aleatoric and epistemic) are output on the screen. On average, the computation takes ∼20 seconds on CPU and ∼10 seconds on GPU. Figure 7a pictorially illustrates the inputs and outputs for predicting the *K_m_* value of a Hexokinase (from *Homo sapiens*) acting on its native substrate D-Glucose. The output value 5.58mM is within 7% error from the experimentally reported value of 6.3mM^48^. In addition, CatPred interface also checks if given inputs already occur in the databases BRENDA and/or SABIO-RK to alert the user. If the check passes, then the database entries corresponding to the inputs are listed (Figure 7b).

**Figure 7.**
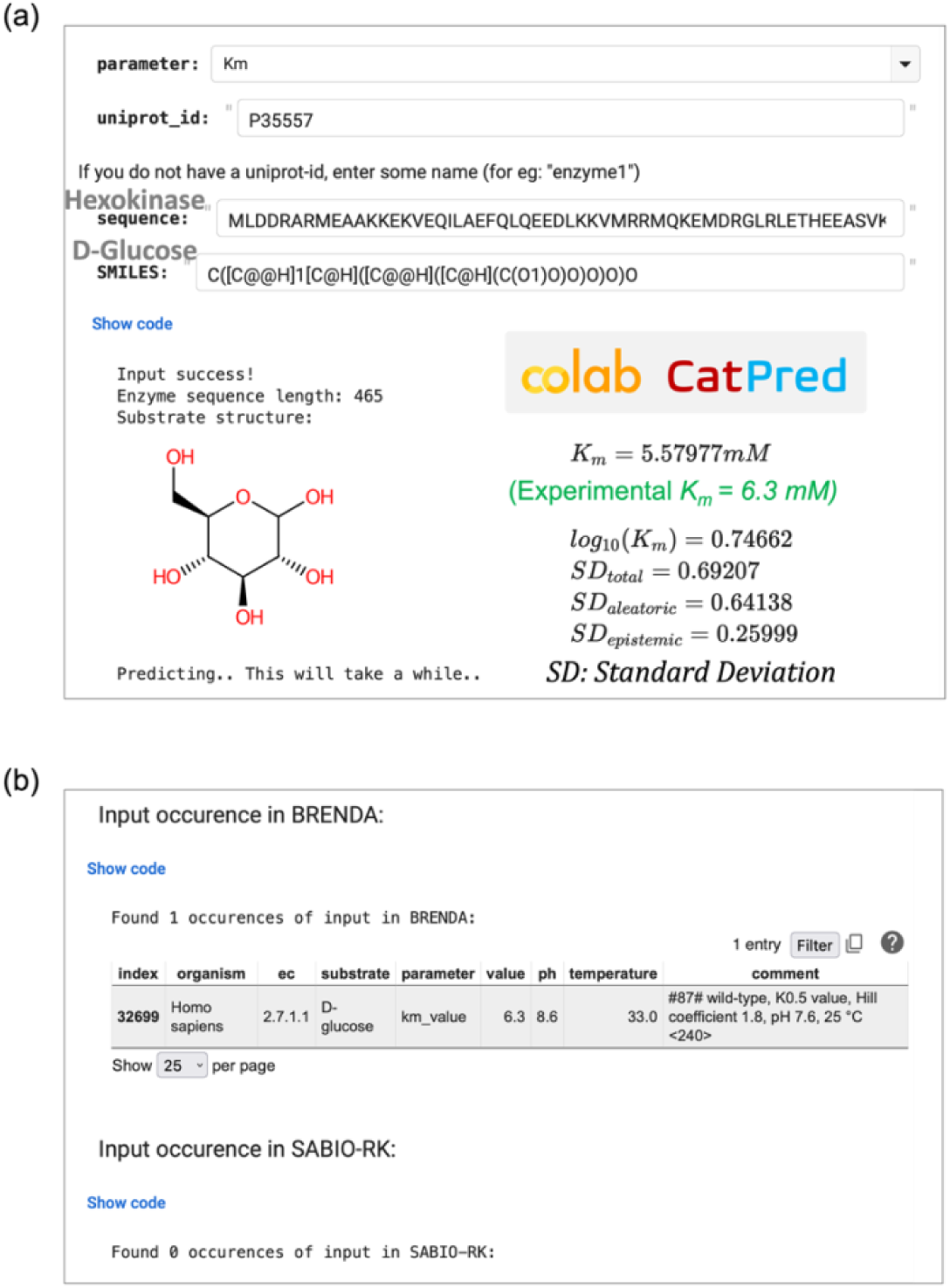
Google Colab interface for making predictions using trained CatPred models. (a) The inputs are the ‘amino acid’ sequence of enzyme and the ‘SMILES’ string of substrate. The predicted output shows the kinetic parameter value (Predicted *Km* value of a Hexokinase enzyme with the D-Glucose substrate in the example shown) and the estimated uncertainty. The contributions to prediction uncertainty (in terms of Standard Deviation: SD in log10-scale) from aleatoric and epistemic uncertainties are also shown. (b) Inputs entered are also searched against entries of the BRENDA and SABIO-RK database. The example input matches with one entry in BRENDA which is shown.

## Discussion

Knowledge of enzyme kinetics is central to the understanding of individual enzymes, metabolic pathways, and dynamic behavior of living cells^24,49,50^. However, experimental determination of enzyme kinetic parameters on a large-scale is an arduous and cost prohibitive task. Although several ML models have been developed before, there is no unified web resource for the prediction of *k_cat_*, *K_m_* and *K_i_* parameters, using standardized training sets, with performance evaluated on out-of-distribution data, and with uncertainty prediction for individual queries. By leveraging rich feature representations and training on expanded and standardized datasets, CatPred achieves performance at least at par with existing studies despite the expanded scope of model coverage.

Prediction quality by CapPred is predominantly limited by the experimental uncertainty in the datasets (i.e., aleatoric uncertainty) as shown in Figure 6. This is confirmed by the fact that ML models trained using different inputs and network architectures arrive at similar metrics of prediction (R^2^ = 0.65 by UniKP^30^ and R^2^ = 0.61 by CatPred for *k_cat_* prediction). Data uncertainty could be ameliorated by directly accounting for environmental conditions such as pH, temperature. Recently, Yu et. al.^30^ trained a *k_cat_* prediction ML model that explicitly considers pH and temperature as inputs and obtained a better accuracy of prediction compared to a baseline model. However, the datasets used for training using pH and temperature were quite small (∼600 datapoints) indicating that these trained models may not be broadly applicable. Such limitations pertaining to datasets call for a systematic effort to generate (and open source) high-quality measurements of enzyme kinetic parameters with complete annotations and broad coverage of enzyme functions. Training on high quality datasets could give rise to model predictions with higher accuracies and lower uncertainties.

We did not find any improvements of prediction performance upon addition of enzyme 3D-structural features extracted using the pretrained E-GNN on top of sequence attention and pLM features. This observation is unsurprising given that the protein language models have previously shown to encode not only sequence but also structural information^51^. Previous works also show that ML models using structural features in addition to pLM features show little improvement over those using pLM features alone^52^. Instead of using entire 3D structures, a targeted description of enzyme-substrate binding regions with information of active-site amino acids could potentially be more informative^53^. Further improvements could focus on incorporating more mechanistic descriptions of enzyme kinetics such as active site and transition state modeling. Different graph neural network architectures can have significant impact on ML model performances. More detailed studies are needed to exhaustively explore these possibilities in context of improving enzyme kinetic parameter prediction.

## Methods

### Dataset curation

The BRENDA database version 2022_2 was downloaded in json format from their website. The SABIO-RK database was downloaded from their website in sbml format. The downloaded databases were processed using in-house Python scripts. All entries of the downloaded databases were parsed while discarding entries that do not have the essential annotations of (1) UniProt identifier for enzyme sequence (or) Organism name and EC number (2) Name of substrate(s) (3) Numerical value of a kinetic parameter (*k_cat_*, *K_m_* or *K_i_*). For entries with a valid Organism name and EC number but no Uniprot-id, Uniprot API search is used to find out all enzyme entries with the given Organism and EC combination. If the search returned a unique enzyme Uniprot-id, the entry was updated with the identified Uniprot-id. Entries belonging to engineered or mutated enzymes were discarded. The Uniprot-identifiers were next used to obtain enzyme sequences and AlphaFold-2.0 predicted structures. Substrate name to SMILES mappings for the entire databases were retrieved from BRENDA and SABIO-RK and used to populate the parsed entries with SMILES strings. For those substrates whose SMILES could not be found on BRENDA and SABIO-RK, we utilized the PubChem’s identifier exchange service (https://pubchem.ncbi.nlm.nih.gov/idexchange/) to obtain SMILES strings. Each SMILES string was canonicalized using the Rdkit Python library. Duplicate measurements (i.e., more than one measurement for the same pair of enzyme sequence and substrate SMILES) were processed by taking the geometric mean of measurements (for *K_m_* and *K_i_*) and the maximum of measurements (for *k_cat_*). This curation process yielded a total of 23,197 *k_cat_*, 41,174 *K_m_* and 11,929 *K_i_* entries with enzyme sequence, enzyme structure, and substrate SMILES. Since the *k_cat_*, *K_m_* and *K_i_* values span several orders of magnitude, the values were log10-transformed to obtain approximately normal distributions for each.

### Dataset splitting

The curated CatPred datasets were split into training (80%), validation (10%) and held-out test sets (10%) using scikit-learn Python package. The splitting ensures that entries in test/validation splits do not have the enzyme sequence and substrate SMILES pairs seen in training splits. The held-out sets are further filtered into subsets based on enzyme sequence identity cutoff to training sequences. Enzyme sequences within each dataset (*k_cat_*, *K_m_* or *K_i_*) are clustered using identity cutoff values of 99%, 80%, 60% and 40% using the mmseqs2^54^ Python library.

### Calculation of enzyme sequence latent spaces

Enzyme sequences were converted into 1280-dimensional numerical representations using the mean features of the final layer of the pretrained ESM-2 model (650 million parameter version). The calculated representations were then clustered into k-nearest neighbor (kNN) graphs with the help of Approximate Nearest Neighbors algorithm^55^ as implemented using the Annoy Python library. The cosine-distance metric was used for clustering. A maximum of 50 kNN trees were built with k value set to 10. Constructed trees were plotted using the TMAP^56^ and Faerun^57^ Python libraries. Two separate plots for CatPred-DB-*k_cat_* and CatPred-DB-*K_m_* were constructed. Within each plot, points were colored according to whether the enzyme sequences newly introduced in CatPred (i.e., were not present in the existing *k_cat_* dataset ^28^ or *K_m_* dataset.^29^) or not.

### Deep learning architecture

The CatPred deep learning framework is built upon that used in ref^58^ and is written in the Python programming language. Each enzyme sequence is first transformed into numerical representation using a neural embedding layer. For CatPred models using Sequence Attention, the sequence embeddings are further enriched with the positional information using Rotary Positional Embeddings^43^ and converted into key, query, values for input to attention layers as described in ref^44^. For CatPred models using protein Language Model features, the ESM2 pretrained model (esm2_t33_650M_UR50D) developed in ref.^7^ is utilized to extract 1280-dimensional features for each enzyme sequence. These features are concatenated with the sequence embedding and attention features. The concatenated features are pooled using an attentive pooling layer that learns a weight for each sequence position and performs a weighted averaging across the sequence length. These pooled features are the final enzyme representations. For each substrate, RDKit is used to generate an atom-bond connectivity graph using the Rdkit Python library. The atoms are converted into features using the corresponding atomic number, number of bonds, formal charge, hybridization, aromaticity, atomic mass, number of hydrogens bonded to the atom and chirality. Each feature is one-hot encoded and concatenated to form the atom feature vector. Similarly, the bonds are converted into features using the bond type (single, double, triple, or aromatic), bond conjugation, bond presence in a ring and bond chirality. These bond features are one-hot encoded and concatenated to form the bond feature vector. The atom and bond features are transformed into molecule features by utilizing the directed-message passing neural network (D-MPNN) as described in ref^41^. Using these, a directed edge feature is constructed for a pair of atoms connected by a bond by concatenating the first atom’s feature with the bond’s features. These edge features are iteratively updated using a learnable neural network with non-linear activation function to aggregate the features of neighborhood atoms^41^. The final molecular representation is obtained by summation of all atom features. The final enzyme and molecular representations are concatenated together and input to a fully connected neural network to output two real values representing the mean and the variance. The E-GNN pre-trained model and its pre-trained weights as described in ref ^45^ are used without any modification to extract the structural features. For each enzyme 3D-structure, this yielded a 128-dimensional embedding. (Supplementary Fig. S3)

### Hyperparameter tuning and training

The hyperparameters of enzyme feature learning modules are: - the dimension of embedding layer, the dimension of rotary positional embeddings, number of attention layers and number of layers in attentive pooling. All hyperparameters of substrate feature learning module were set to optimal values recommended in ref^58^. The learning rate was fixed at 0.001 and the batch size was tuned accordingly. The number of models in the ensemble when training CatPred models was set to 10. The rectified linear unit (relu) activation function was used for all layers except for the output layers. All the models were trained in batches using the Adam optimizer and the training dataset was fed into the model for 20 epochs. We used minimization of the negative log-likelihood loss function as the objective function as described in ref^39^. Different combinations of listed hyperparameters were tried to train models and optimal values are chosen by the performance of trained models on the validation dataset. The optimal values so obtained are used to train models on the training+validation and training+validation+test datasets for testing and production purposes respectively. The production models were trained for 30 epochs. The list of tested hyperparameters and the obtained optimal values are listed in a detailed architecture block figure Supplementary Fig. S3.

## Data availability

CatPred-DB datasets will be made publicly available upon publication at https://github.com/maranasgroup/catpred

## Code availability

All the codes corresponding to the experiments presented in the manuscript will be made publicly available upon publication at https://github.com/maranasgroup/catpred

A web interface for using trained CatPred models is currently available at https://tiny.cc/catpred

## Acknowledgements

This work was funded by the DOE Center for Advanced Bioenergy and Bioproducts Innovation (U.S. Department of Energy, Office of Science, Office of Biological and Environmental Research under Award Number DE-SC0018420). Funding also provided by the Center for Bioenergy Innovation, which is a U.S. Department of Energy Bioenergy Research Center supported by the Office of Biological and Environmental Research in the DOE Office of Science. Oak Ridge National Laboratory is managed by UT-Battelle, LLC for the US DOE under Contract Number DE-AC05-00OR22725. This material is based upon work supported by the Center for Bioenergy Innovation (CBI), U.S. Department of Energy, Office of Science, Biological and Environmental Research Program under Award Number ERKP886. Any opinions, findings, and conclusions or recommendations expressed in this publication are those of the author(s) and do not necessarily reflect the views of the U.S. Department of Energy.

Funding was also provided by the DOE Office of Science, Office of Biological and Environmental Research (Award Number DE-SC0018260). This work was also supported by the U.S. National Science Foundation funded Molecule Maker Lab Institute (MMLI), award number 2019897 supported by National AI Research Institutes Program of the Directorate for Computer and Information Science and Engineering (CISE), in collaboration with the Division of Chemistry (CHE) and the Division of Chemical, Bioengineering, and Environmental Transport Systems (CBET) awarded to CDM. The funders had no role in study design, data collection and analysis, decision to publish, or preparation of the manuscript.

## Notes

https://github.com/maranasgroup/CatPred

https://tiny.cc/catpred

### Competing Interest Statement

The authors have declared no competing interest.

### Summary of Updates

Added results of EGNN models to figures and text. Revised figures for clarity and legibility. Some fonts were missing in the figures within the biorxiv generated pdf. This has been fixed. Supplementary files have been updated accordingly as well.

