## Supplementary Information for "CatPred: A comprehensive framework for deep learning in vitro enzyme kinetic parameters *k_cat_*, *K_m_* and *K_i_*"

**Table S1. Statistics of *in vitro* kinetic measurements in BRENDA (v2022_2) and SABIO-RK (as of Nov. 2023)**

|  | ***k_cat_*** | ***K_m_*** | ***K_i_*** |
| --- | --- | --- | --- |
| **BRENDA** | 83,662 | 172,917 | 45,684 |
| **SABIO-RK** | 16,956 | 37,447 | 8,179 |

**Table S2. Number of unique Enzyme Commission (EC) for BRENDA (v2022_2) entries with kinetic parameters**

|  | ***k_cat_*** | ***K_m_*** | ***K_i_*** |
| --- | --- | --- | --- |
| **Unique EC numbers** | 3,336 | 4,885 | 1,986 |

**Table S3. CatPred-DB dataset proportions with annotations of ‘natural substrate’**

|  | ***CatPred-DB-k_cat_*** | ***CatPred-DB-K_m_*** | ***CatPred-DB-K_i_*** |
| --- | --- | --- | --- |
| **Natural** | 11,107 (47.8%) | 23,989 (58.3%) | 3,304 (27.7%) |
| **Total** | 23,197 | 41,174 | 11,929 |

**Table S4. CatPred hyperparameter list – choices tested, and final values chosen. All other hyperparameters were not tuned and set to recommended values in chemprop**^1^ **and are available on our Github repository.**

|  | ***Choices tested*** | ***Final value chosen*** |
| --- | --- | --- |
| **Batch size** | [32, 64, 128] | 32 |
| **Sequence Embedding size** | [36, 64, 128, 360, 480] | 36 |
| **Attention heads** | [3, 6, 12] | 6 |

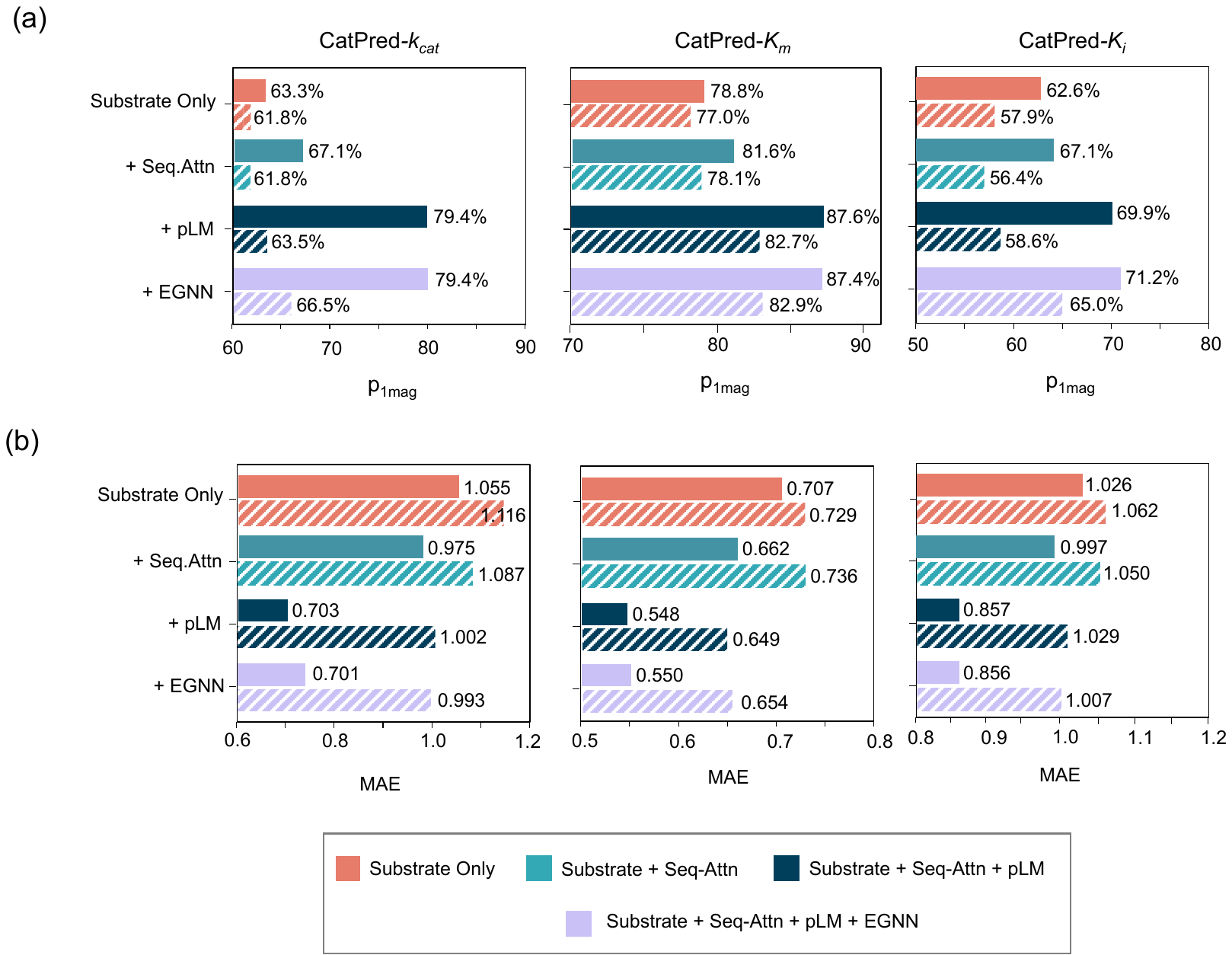

**Figure S1.** The performance metrics achieved by CatPred-*k_cat_*, CatPred-*K_m_* and CatPred-*K_i_* models on hold-out test sets (solid bars) and on out-of-distribution samples (patterned bars). The (a) Percent of predictions within one order of magnitude error (p_1mag_) and (b) Mean Absolute Error (MAE) are shown. ‘Substrate Only’ refers to CatPred models trained using only the substrate features; ‘Substrate+Seq-Attn’ (Sequence Attention) refers to CatPred models trained using substrate features and the Seq-Attn features; ‘Substrate+Seq-Attn+pLM’ (protein Langugae Model) refers to CatPred models trained using substrate features along with both the Seq-Attn and pLM features. ‘Substrate+Seq-Attn+pLM+EGNN’ (Equivariant Graph Neural Networks) refers to CatPred models trained using substrate features along with Seq-Attn+pLM and EGNN features.

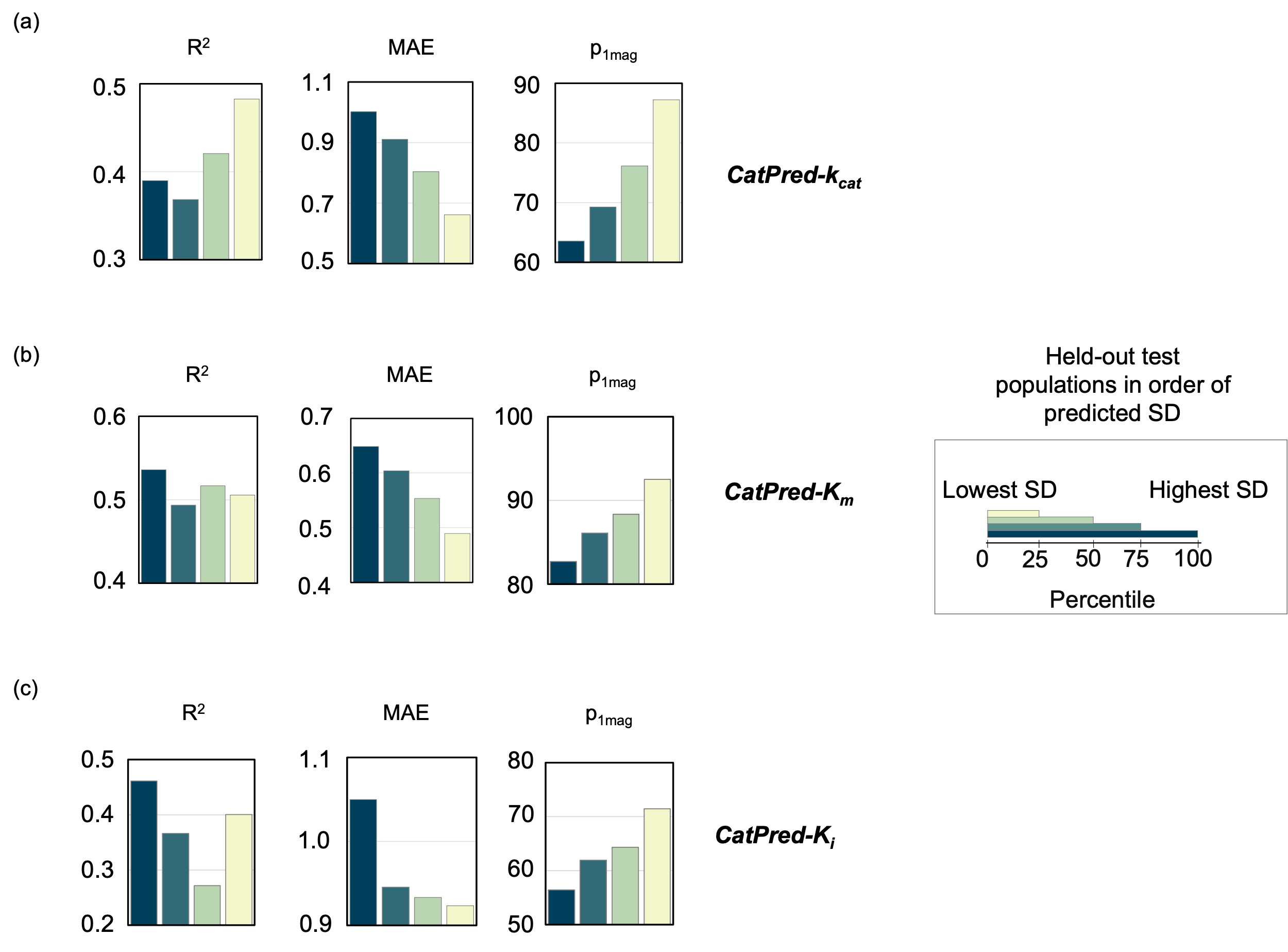

**Figure S2** The performance metrics achieved by (a) CatPred-k_cat_, (b) CatPred-K_m_ and (c) CatPred-K_i_ models on sub populations of the out-of-distribution test sets binned in order of their predicted uncertainty values (sum of aleatoric and epistemic uncertainty). Each colored bar denotes a sub population of the held-out set with uncertainty less than the 100^th^ (Blue), 75^th^ (Dark Green), 50^th^ (Light Green), and 25^th^ (Light yellow) percentile respectively. Within each figure, the subplots show the obtained co-efficient of regression (R^2^), Mean Absolute Error (MAE) and Percent of predictions within one oder of magintude error (p_1mag_).
